# Retroviral origins of the *Caenorhabditis elegans* orphan gene F58H7.5

**DOI:** 10.1101/073510

**Authors:** Wadim J Kapulkin

## Abstract

This work describes the results of the genome-scale analysis of endogenous retrovirus insertions in two *C. elegans* isolates: the prototype N2 (Bristol) and CB4856 (Hawaii). In total thirteen, identification of potentially replication competent, endogenous retroviral elements is described. Ten elements were identified as conserved between N2 and CB4856 by the reciprocal match of paired LTRs. The description focuses on the particular endogenous retrovirus insertion wich is identified on the proximal arm of the chromosome IV (located at positions IV: 912,948 – 921,658 and IV: 899,767 – 908,485 of the N2 and CB4856 respectively). In both isolates the inserted provirus is flanked by the predicted long terminal repeats (LTR)s of the length of 415 bp and of identical sequence. Provided the absolute LTR sequence identity this particular provirus represents insertion acquired prior to split from the common ancestor, suggesting this insertion event is evolutionary recent. The identified insertion of the endogenous retrovirus embeds the orphan gene F58H7.5, specific to *C. elegans* lineage. This unprecedented example establishes that in the evolutionary past *C. elegans*, had acquired the gene of the retroviral origins presumably via mechanisms involving the RNA intermediate.

**Importance:** This work describes the retroviral origins of *C. elegans* orphan gene F58H7.5. Presented work implies that in the evolutionary past the *C. elegans* have acquired new gene as a result of the infection event. *C. elegans* is presently regarded as genetic model organism widely used in genetic research. The genome of *C. elegans* have been sequenced nearly 20 years ago. This unprecedented example establishes that in the evolutionary past C. elegans genome, had acquired the gene of the retroviral origins presumably via mechanisms involving the RNA intermediate.

## Introduction

Endogenous retroviruses and LTR retrotransposons are ubiquitously present in eukaryotic genomes. Endogenous retroviruses and LTR retrotransposons are widely assumed to replicate via RNA intermediates. Endogenous retrovirus insertions present in the genomes are thought to represent the remnants of the infectious events of the past evolutionary history. Examples of an active endogenous retroviruses however are found almost invariably among metazoan genomes, and contribute to the reverse flow of the genetic information. Benign endogenous retroviruses that are found inserted into genomes of animals and are known to contribute to phenotypic traits and overall genetic variation. Importantly, certain endogenous retroviruses are known for the potential to convert into active retroviruses (i.e. Moloney Leukemia Virus (Stoye and Coffin 1987)) and Mouse Mammary Tumor Virus (Bittner 1936)) and those are grouped together with circulating retroviruses (i.e. Rous Sarcoma Virus (Rous 1911)) often with significant pathogenic potential an therefore pose an active and significant threat to the health of the animal populations. Infectious properties of the retroviral particles led to development of the retroviral vectors as means to efficiently insert proviral DNA into genomes. In contrast, particularly well studied genetic model organism, *C. elegans* appears to harbor considerably fewer LTR elements than other animal models. Further, sparse endogenous retroviruses inserted into *C. elegans* appear mostly dormant and do not appear to significantly contribute to observable traits. One notable exception is the RETR-1 element (Britten 1995) inserted into *C. elegans* plg-1 locus, (Papoli et al. 2007) contributing to natural variation in the copulatory plug polymorphism (Doniach and Hodgkin 1995). The active transposition and replication of the RETR-1 however has not been demonstrated (Preiss 2007), neither any of the other endogenous (nor exogenous) retroviral elements present in *C. elegans* genome. This work describes the identification of the ‘first –pass set’ of the potentially replication competent proviruses inserted into *C. elegans* genome present in both N2 (Bristol) and CB4856 (Hawaii) lines. The description is focused on the distinct and unique proviral insertion on the proximal arm of the chromosome IV identified by the reciprocal LTR match. This particular prediction distinctly embeds the orphan gene of *C.elegans* F58H7.5. Given the recent improvement in the genome engineering i.e. with the advancement in the CRISPR-Cas9 methods (Jinek et al. 2012) applicable to *C.elegans* (Friedland et al. 2013), I suggest the identified set might serve the practical and experimental purpose: the proviruses once inserted into the *C.elegans* genome in the evolutionary past could now be ablated (Yang et al. 2015).

## Materials and methods

Genebank-NCBI reference sequence of the prototype N2 (Bristol) *C. elegans* genome (C. elegans Sequencing Consortium 1998): Chr I NC_003279.8, Chr II NC_003280.10, Chr III NC_003281.10, Chr IV NC_003282.8, Chr V NC_003283.11, Chr X NC_003284.9 and Hawaiian isolate CB4856 (Thompson et al. 2015): I Chromosome CM003206.1, II Chromosome CM003207.1, III Chromosome CM003208.1, IV Chromosome CM003209.1, V Chromosome CM003210.1, X Chromosome CM003211.1.

*C. elegans* chromosome sequence assemblies were analyzed with LTR finder program (Zhao Xu, Hao Wang 2007) at default parameters with tRNA primer binding site predicted using *C. elegans* tRNA dataset, to detect individual flanking LTR pairs. The first-pass screening, focused on the predictions were at least one internal candidate reading frame were suggested by internally enabled ScanProsite. DNA sequences predicted by the LTR finder were conceptually translated into reading frames (<molbiol.ru/eng/scripts/01_13.html>) resulting in strings of the one-letter amino-acid codes. Individual reading frames of coded amino-acid strings were masked for in-frame stop codons. Stop-masked amino-acid strings were analyzed with the external InterProScan search engine (<www.ebi.ac.uk/Tools/pfa/iprscan/> Zdobnov et al. 2001) for the retroviral domain detection and output was cross-verified with the output of the LTR finder. LTR identity in an isolate matched candidate predictions of the LTR pairs were verified by the reciprocal sequence alignment (Altschul et al. 1990). Predicted reading frames were verified with analysis by taxon restricted homology search (either with the set of the NCBI reference sequence of the retro-transcribing viruses or endogenous retroviruses). Identified proviruses were displayed on Ensembl *C. elegans* N2(Bristol) database by BLAT (Kent 2002). The protein domains architectures ideograms and the pairwise LOGO alignments were drawn with MyDomains prosite (Hulo et al. 2008) (<prosite.expasy.org/mydomains/>) and Weblogo3 (Crooks ey al. 2004, Schneider and Stephens 1990) (<weblogo.threeplusone.com>) respectively. The genomic landscape surrounding the confirmed retroviral insertions was GBrowse (<gbrowse.org>) displayed at the WormBase website. Phylogenetic analysis on predicted retroviral domains was conducted with pipeline in ETE3 at (<www.genome.jp>). Alignment and phylogenetic reconstructions were performed using the function “build” of ETE3 v3.0.0b32 (Huerta-Cepas et al., 2016) as implemented on the GenomeNet (<www.genome.jp/tools/ete/>). The compiled input files for reverse-transciptase polymerase, RNaseH and integrase catalytic domain are given in the supplement files. Alignment was performed with MAFFT v6.861b with the default options (Katoh and Standley, 2013). The initial tree was constructed using FastTree v2.1.8 with default parameters (Price et al., 2009). The ML bootstrapped trees were inferred either using RAxML v8.1.20 ran with model PROTGAMMAJTT and default parameters (Stamatakis, 2014) and PhyML v20160115 ran with model JTT and parameters: -f m --pinv e -o tlr --nclasses 4 --bootstrap 100 --alpha e (Guindon et al., 2010). Branch supports were computed out of 100 iterations.

## Results

The details of the genome-scale analysis are included in materials and methods section and the results of the first pass screening are listed with (Table 1. and Table 4.) and graphically outlined (Fig. 5), however, will be described elsewhere. Here, I describe the identification and analysis of the unusual provirus inserted in sense (+) orientation on Chromosome IV of both prototype N2(Bristol) strain and Hawaiian isolate CB4856. The provirus is found inserted at the proximal arm of IV at positions IV:912948..921658 and IV:899767..908485 of an N2 and CB4856 respectively by the reciprocal match of the LTR pairs (Fig. 1.A. and B). The two extracted sequences (8711 bp and 8719 bp respectively) align precisely to ensembl N2(Bristol) assembly, with duplicate matches corresponding to long terminal repeats (Fig. 1. C.). This suggests the two elements identified at the proximal arm of the chromosome IV, might represent provirus inserted in the ancestral *C. elegans* lineage, which have independently diverged since the insertion time. To prove the identified provirus share a common ancestor, 1kb of immediately flanking the insertion sequence was extracted and aligned, confirming the 5’ and 3’ flanks were almost identical and have diverged independently by four base substitution per thousand nucleotides (0.4%) (Supplement file.1. Alignment of the insertions with 1kb flanking sequence). The striking sequence conservation detected in the provirus flanking sequence between N2(Bristol) and CB4856, indicates the element once inserted into the genome of the ancestral *C. elegans* line, was inherited as a haplotypic block and presumably maintained in same location on chromosome IV in both isolates.

**Figure 1.**
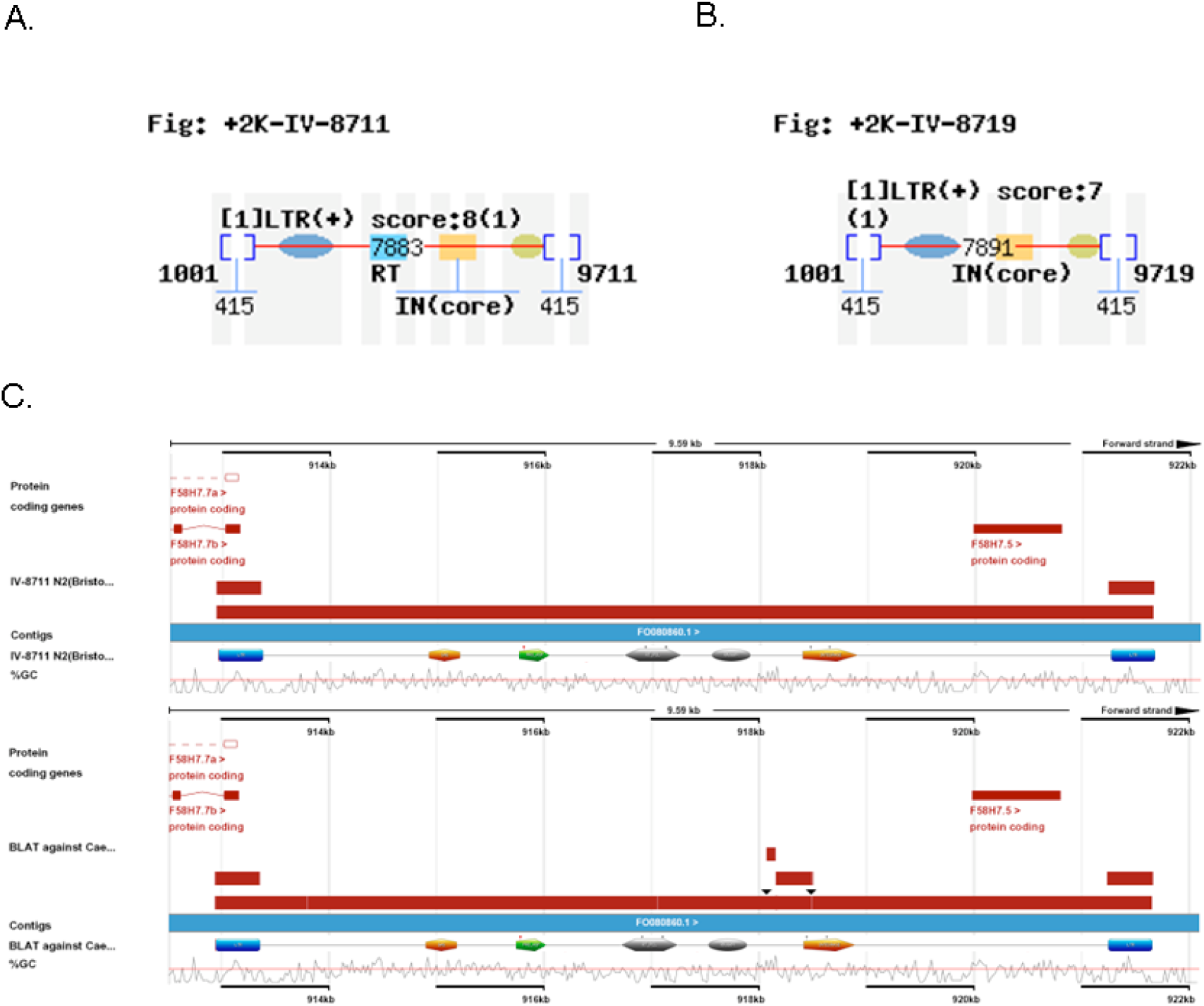
LTR finder graphics representing predicted provirus on the proximal arm of the chromosome IV in N2 (A.), and CB4658 (B.) Blue oval represents conserved PBS and yellow oval represents conserved PPT. Note RT and Integrase domains are predicted in N2 (A.) but only integrase domain is detected in CB4856 (B.). Ensembl representation (C.) of BLAT alignment of provirus sequences N2 (upper panel), and CB4658 (lower panel) with imposed Prosite Domain architectures (415 bp LTRs are drawn as blue rectangles) scaled onto Ensembl BLAT images (above the track with %GC content). The red horizontal bars represent BLAT alignments, expectedly duplicated at the 5’ and 3’ LTR regions (upper panel). Additional BLAT match is apparent with IV-8711 provirus of an N2 aligned with chromosome IV assembly of CB4856 (lower panel). The extra bar represents non-continuous alignments in the RNaseH – integrase connecting region (indicated by black arrowheads). Note the 3’ region of the non-continuous alignment overlaps with N-terminal region of the predicted integrase (see Fig. 2 for details). F58H7.5 is apparent in both panels in the sector describing protein coding genes annotated in the reference sequence assembly. Provided the ideograms are drawn up to scale F58H7.5 projects on the 3’ sub-LTR region of the predicted provirus.

**Figure 5.**
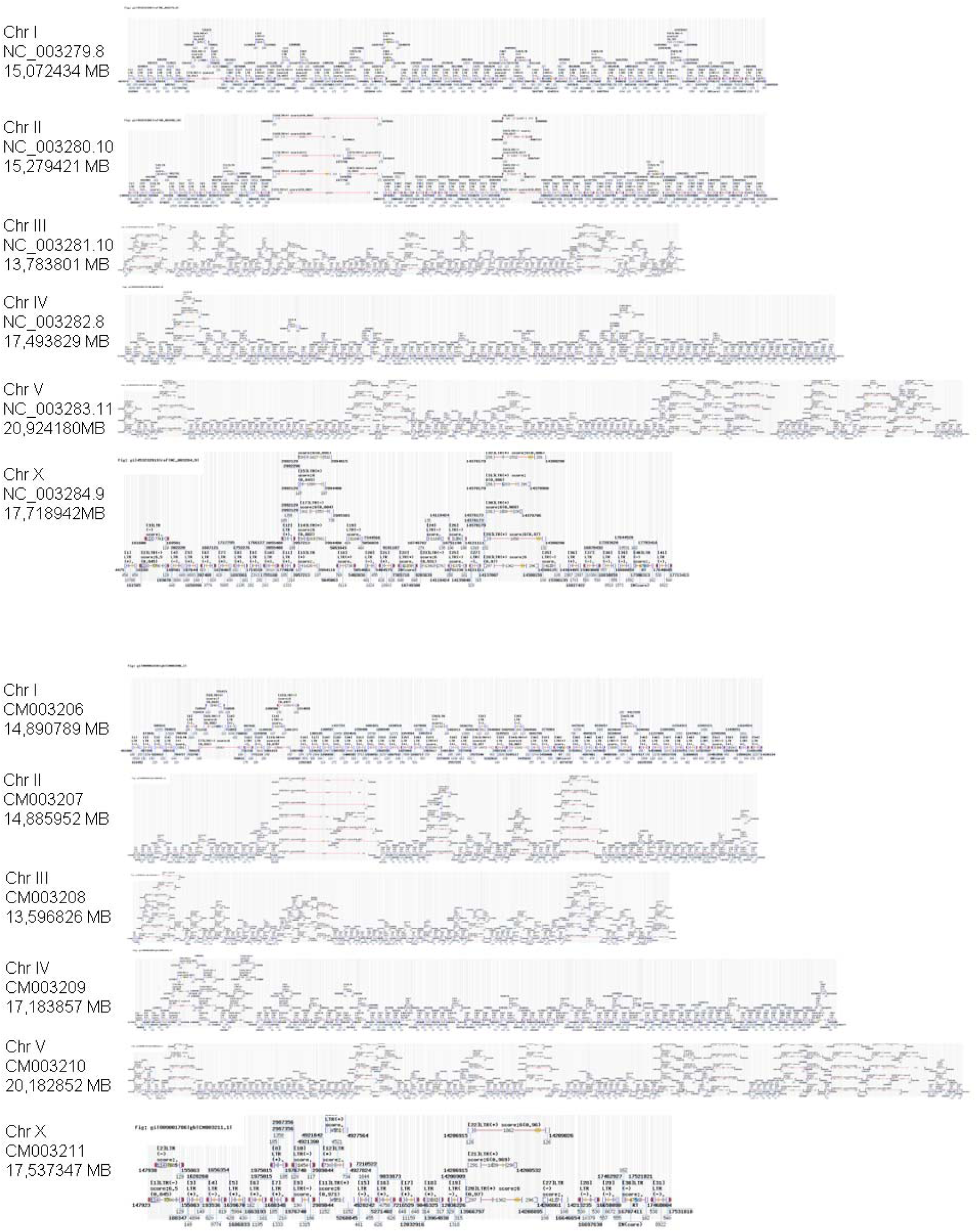
Genome-scale LTR finder predictions of the candidate LTR pairs. A. Chromosomal representation of the LTR finder output pn N2(Bristol) genome, B. Chromosomal representation of the LTR finder output on CB4856(Hawaii). The description is included into Tables 1. and 4.

**Table 1.** Results of the first-pass genome-scale LTR finder predictions of the endogenous retroviral elements inserted in chromosomal assemblies of two C. elegans isolates: reference N2(Bristol) and CB4856(Hawaii).

| Chr | Retroviral insertion LTR name | N2 insert length | Hawaii insert length | N2 chromosome position | N2 LTR 5'/3' | Hawaii chromosome position | Hawaii LTR 5'/3' |
| --- | --- | --- | --- | --- | --- | --- | --- |
| I | I-CeN2ii-1879 (*) | 1879 | 1879 | 13147285 - 13149163 | 245/<br>245 | 12993371 – 12995249 | 245/<br>245 |
|  | I-CeN2ii-5088 | 5088 | 5088 | 3847365 - 3852452 | 217/<br>221 | 3776964 - 3782051 | 217/<br>221 |
| II | II-CeN2-16388 | 16388 | - | 13243641 - 13260028 | 161/<br>161 | - | - |
| III | III-CeN2-8886 (RETR-1) | 8886 | - | 8852597 - 8861482 | 518/<br>513 | - | - |
|  | III-CeN2ii-13588 | 13588 | 13506 | 13229612 - 13243199 | 517/<br>517 | 13044879 - 13058384 | 517/<br>517 |
| IV | IV-CeN2ii-8711 | 8711 | 8719 | 912948 - 921658 | 415/<br>415 | 899767 - 908485 | 415/<br>415 |
| V | V-CeN2ii-10045 | 10045 | 10048 | 4300964 - 4311008 | 318/<br>318 | 4041012 - 4051059 | 316/<br>320 |
|  | V-CeN2-16124 | 16124 | - | 8473016 - 8489139 | 180/<br>193 | - | - |
|  | V-CeN2ii-19860 | 19860 | 19794 | 8825452 - 8845311 | 514/<br>514 | 8520474 - 8540267 | 514/<br>514 |
|  | V-CeN2-11685 | 11685 | - | 17146405 - 17158089 | 444/<br>457 | - | - |
|  | V-CeN2ii-19417 | 19417 | 19417 | 18435912 - 18455328 | 251/<br>244 | 18435912 - 18455328 | 251/<br>244 |
|  | V-CeN2ii-12067 | 12067 | 12540 | 19270045 - 19282111 | 537/<br>537 | 18647618 - 18660157 | 537/<br>537 |
| X | X-CeN2ii-12453 (**) | 12453 | 12453 | 9191187 - 9203639 | 648/<br>648 | 9033873 - 9046325 | 648/<br>648 |
|  | X-CeN2ii-5078 | 5078 | 5078 | 17644528 - 17649605 | 162/<br>162 | 17462927 - 17468004 | 162/<br>162 |

(\*) - denotes unusually short element. (\*\*) – denotes insertion identified on homologous CB4856 chromosome by the reciprocal LTR finder match only.

**Table 4.**
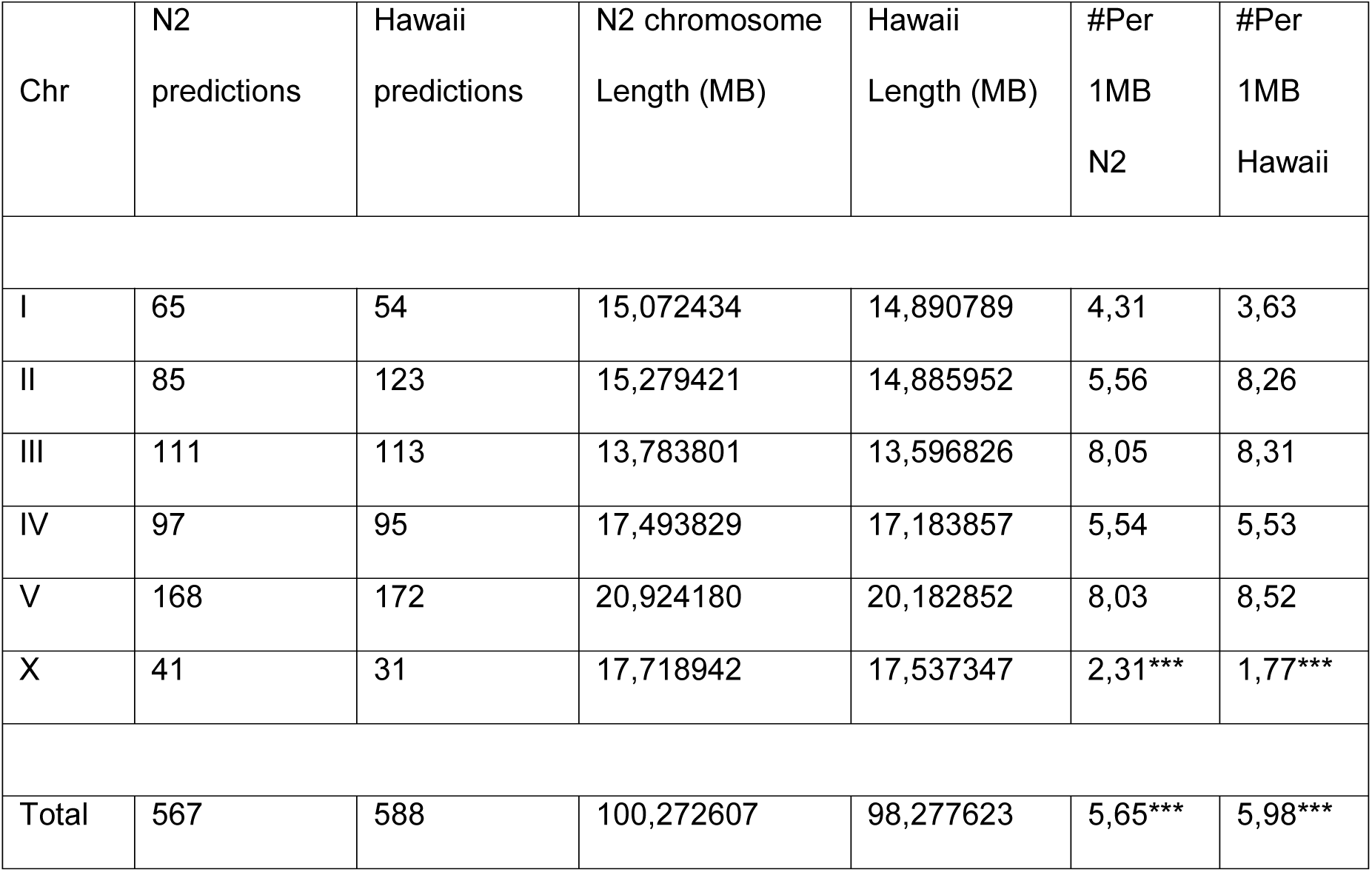
LTR finder calculations on the number of the genome scale predictions per chromosome. Values do vary, ranging from 2,31 - 1,77 (for chromosome X) to 8,05 and 8,52 (for chromosome III and chromosome V) LTR finder predictions per MB in N2(Bristol) and CB4856(Hawaii) respectively. While the lowest number of LTR finder predictions per MB is observed consistently for sex chromosomes in both isolates, when autosomal predictions are included into calculations the association between numbers of predictions calculated on two genomes are considered extremely statistically significant (*** two tailed P<0.0001, Chi-square test). The reason for *C. elegans* X chromosomes scoring the lowest values remains to be determined, but likely reflects the proportionally lower number of the haplotypic duplications maintained on sex chromosomes when compared to autosomes.

| Chr | N2<br>predictions | Hawaii<br>predictions | N2 chromosome<br>Length (MB) | Hawaii<br>Length (MB) | #Per<br>1MB<br>N2 | #Per<br>1MB<br>Hawaii |
| --- | --- | --- | --- | --- | --- | --- |
| I | 65 | 54 | 15,072434 | 14,890789 | 4,31 | 3,63 |
| II | 85 | 123 | 15,279421 | 14,885952 | 5,56 | 8,26 |
| III | 111 | 113 | 13,783801 | 13,596826 | 8,05 | 8,31 |
| IV | 97 | 95 | 17,493829 | 17,183857 | 5,54 | 5,53 |
| V | 168 | 172 | 20,924180 | 20,182852 | 8,03 | 8,52 |
| X | 41 | 31 | 17,718942 | 17,537347 | 2,31*** | 1,77*** |
| Total | 567 | 588 | 100,272607 | 98,277623 | 5,65*** | 5,98*** |

## LTRs - Long Terminal Repeats

(Fig 1.A. and B. and supplement files). The LTRs are identical in length in both *C.elegans* isolates. N2(Bristol) 5'-LTR is located at IV:912,948 – 913,362 and 3'-LTR IV:921,244 – 921,658 and both LTRs are of same length 415 bp. Hawaii CB4856 5'-LTR is located on IV:899,767 – 900,181 and 3'-LTR IV:908071 – 908485 and both LTRs are of same length 415 bp (Supplement file 2. LTR alignment). Comparison between 5’ LTRs and 3’ LTRs in both isolates confirms the lack of divergent bases in the LTR alignment. Provided the accepted model of the retroviral replication (Telesnitsky A and Goff SP 1998) assumes the LTRs are identical at the insertion time, therefore the lack of divergence in the LTR sequences in both geographical isolates suggests that the identified insertion represent the recent evolutionary event.

## PBS

Primer Binding Sites (Fig 1.A. and B. and supplement files) are predicted by LTR finder as sequences immediately downstream (12nt) of 5’ LTR matching the tRNA primer for the reverse transcription of viral RNA (Telesnitsky A and Goff SP 1998). The LTR finder program identified the same iso-accepting methionyl (CAT) tRNA to bind conserved PBS [5’-TAGCTAGCGAGTGAACCGAATTTCG] (IV:913,374 – 913,398) in the C.elegans N2(Bristol) insertion and Hawaiian CB4856 isolate (IV:900,193 – 900,217).

## PPT

Poly Purine Tract sequence (Fig 1.A. and B. and supplement files) immediately proceeding the 3’ LTR are identified by the LTR finder in both proviruses and are identical in two analyzed isolates [PPT: 5’-TCAAAAGGGGGGAGG] located at (IV:921,229 – 921,243) in the C.elegans N2(Bristol) insertion and (IV:908,056 – 908,070) Hawaiian CB4856 isolate.

The insertions of the provirus at the proximal arm of the chromosome IV, are almost identical in the overall length. The provirus in the reference N2(Bristol) assembly is 8,711 nt long and CB4856(Hawaii) is minimally longer 8, 719 nt (Table 1. and supplement file). Given the LTRs, are of identical length in both isolates and essentially lack any aberrant bases, the region between LTRs (coding for retroviral proteins) is expected to harbor divergent nucleotides accounting for the difference in the overall length. BLAT analysis indicates the two regions (Fig.1 C. indicated by the black arrowheads) interrupting the concordance of the alignment of the provirus variant found in chromosome IV in CB4856 and the reference N2 assembly. Those two regions are displayed in pairwise alignment (Supplement file 1.) and occur in region of provirus at bases in range 5,00-5,57kb (Fig. 2) in a region connecting segments encoding for reverse transcriptase-RNaseH and catalytic integrase proteins. Compared regions in the reference assembly of an N2 and CB4856 assembly, identifies the following segments as inconsistent between isolates: 1. Four base deletion in N2 (deletion flank GAC TTC----CGA CGC) or (ATGT) insertion in CB4856. 2. Three base insertion in N2 (insertion flank CTG GGT – TCG CCG) or (TGC) deletion in CB4856. 3. Seven base pair deletion in N2 (deletion flank AGT AGT ACA TGG) or (ATCATGA) insertion in CB4856, with G to T single base substitution in the left flank region. Collectively across the above 5,00-5,57kb region cumulative aberrant bases (including two substitutions not altering the length) contribute to the overall ~2,8% of divergence between two isolates.

**Figure 2.**
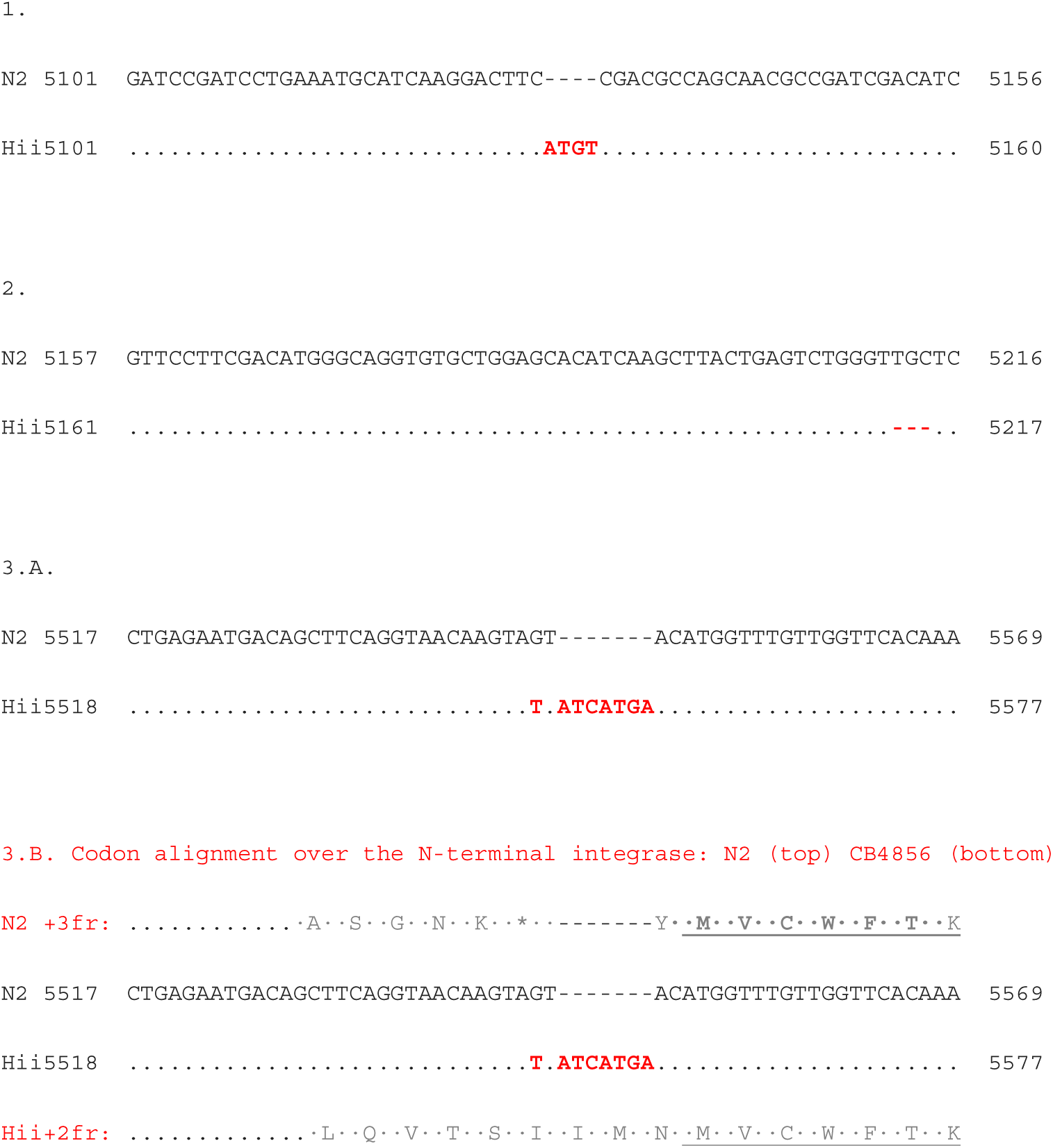
Insertions and / or deletions in the region connecting the RNaseH and catalytic integrase domain encoding region of IV-8711 in N2(Bristol) and IV-8719 in CB4856(Hawaii). The frame altering (3.A.) lesion overlaps with region encoding for the predicted N-terminal catalytic integrase domain (outlined in Fig.1.D.). Residues corresponding to N-terminal catalytic integrase domain alignment start are underlined (3.B). Note the non-synonymous substitution (T->G) in N2 sequence introduces the amber stop codon.

As indicated in (Fig.1.A and 1.B), the provirus inserted into N2(Bristol) is assigned two retroviral domains (Reverse transcriptase RT-domain: at IV:916,714 – 917,250 and Integrase catalytic domain: IV:918,494 – 918,889) by the LTR finder internally enabled PrositeScan. In contrast, CB4856 inserted element is attributed with only one retroviral domain (Integrase catalytic domain: IV:905,294 – 905,716). Provided high level of the overall sequence conservation over the entire length of the insertion present in both isolates, that asymmetric display is somewhat unusual for the elements shared between N2 and CB4856. To address this discrepancy, the proviruses identified in N2 and CB4856, were subjected to analysis with external protein domain search the InterProScan.

## Predicted proviral proteins and protein domains

Provirus IV-8711 in N2(Bristol) and IV-8719 in CB4856(Hawaii) exhibits organization of protein domains typical for endogenous retroviruses/retrotransposons (Fig.1.C). The protein domains predicted by the external InterProScan are presented as pairwise consensus LOGO alignment of particular N2/CB4856 protein domain pairs in (Fig.3.). The protein domains predicted by the external InterProScan sequence LOGO’s are presented in 5’->3’ order as they occur encoded in the provirus (Fig.3.A, gag; B. protease; C. RT polymerase; D. RNase H; E. integrase) with respect to predicted reading frames. The summarized InterProScan search results (below) demonstrate that the provirus identified in IV-8711 in N2(Bristol) and IV-8719 in CB4856(Hawaii) in overall genetic organization appears typical for the endogenous retroviruses. However an additional open reading frame is found in the IV-8711/IV-8719 provirus, encoded by the ORF present in the region between gene encoding the catalytic integrase protein and the 3’-LTR. This sub-3’-LTR localization is typical for the retroviral envelope protein (env), but provided the lack of the homology with retroviral env genes is termed below 3’-ORF and represent the uncharacterized *C. elegans* protein F58H7.5 (Fig 1.C).

**Figure. 3.**
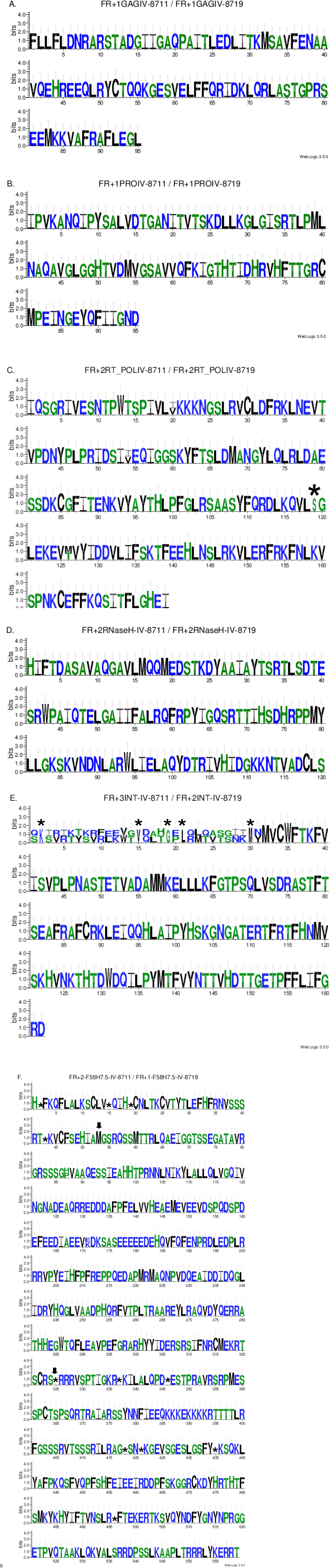
Frame-wise LOGO alignments of the predicted protein domains on proviral insertions on the proximal arm of the chromosome IV present in two *C. elegans* isolates: IV-8711 N2(Bristol) and IV-8719(Hawaii). Group specific antigen gene (gag) (A.); Aspartyl protease (pro) (B.); Reverse-transcriptase polymerase (pol) (C.); RnaseH (D.); Catalytic integrase (int)(E.); 3’-ORF {F58H7.5} (F.). Stop codons are indicated by (*). Black arrows indicate the start and end of the F58H7.5 coding region embedded into the 3’-ORF sequence.

i. gag. Protein domain specifying the gag gene is defined by the Pfam protein domain signature match (PF03732/InterPro:IPR005162). Gag is encoded in the reading frame (+1) and is identical between IV-8711 in N2(Bristol) and IV-8719 in CB4856(Hawaii) proviruses (Fig.3.A). Protein domain encoding for gag is 95 amino-acid residues long and encoded in positions: (1951-2235) in both isolates.
ii. pro. Protein domain specifying the Retroviral-type Aspartyl Protease gene is defined by the Pfam protein domain signature match (PF13650). Aspartyl protease is encoded in the reading frame (+1) and is identical between IV-8711 in N2(Bristol) and IV-8719 in CB4856(Hawaii) proviruses (Fig.3.B). Protein domain encoding for aspartyl protease is 95 amino-acid residues long and encoded in positions: (2791-3075) in both isolates.
iii. pol. Protein domain specifying the Reverse-transcriptase polymerase gene is defined by the ProSite protein domain signature match (PS50878/InterPro:IPR000477). Reverse-transcriptase polymerase is encoded in the reading frame (+2) and is not identical between IV-8711 in N2(Bristol) and IV-8719 in CB4856(Hawaii) proviruses (Fig.3.C). Protein domain encoding for aspartyl protease is 179 amino-acid residues long and encoded in positions: (3767-4303) in both isolates. Across the Reverse-transcriptase polymerase sequence there are three non-synonymous substitutions detected (positions 20 and 54 altering I->V and position 126 M->T) diverged IV-8711 in N2(Bristol) and IV-8719 in CB4856(Hawaii) proviruses. In addition, the reference N2 sequence contains in-frame stop codon predicted in position 119, absent in CB4856, substituting for the serine in this position (Fig.3.C).
iv. RNaseH. Protein domain specifying the Retroviral RNaseH gene is defined by the CDD protein domain signature match (cd09274). RNaseH is encoded in the reading frame (+2) and is identical between IV-8711 in N2(Bristol) and IV-8719 in CB4856(Hawaii) proviruses (Fig.3.D). RNaseH protein domain is 121 amino-acid residues long and is encoded in positions: (4583-4954) in both isolates.
v. int. Integrase catalytic domain. Protein domain specifying the catalytic integrase gene is defined by the ProSite protein domain signature match (PS50994/InterPro:IPR001584). Markedly the Integrase is encoded in the different reading frames: (+3) and (+2) frames are used in IV-8711 in N2(Bristol) and IV-8719 in CB4856(Hawaii) proviruses respectively (Fig.3.E) and (Table. 2). This frame-shift is due to mutations listed in (Fig.2.3) and explains the lack of an alignment in N-terminal part of predicted integrase sequence, until the metionine residue in the position 32. Remaining part of the protein domain encoding for integrase is 130 amino-acid residues long and is identical between IV-8711 in N2(Bristol) and IV-8719 in CB4856(Hawaii) proviruses. Integrase catalytic domain is encoded in positions: (5457-5942) and (5465-5950) in IV-8711 in N2(Bristol) and in IV-8719 in CB4856(Hawaii) respectively.
vi. 3’-ORF. 3’-ORF embeds an uncharacterized protein F58H7.5 predicted in chromosome IV of the reference sequence assembly of the *C. elegans* genome (Fig. 1. C). In the InterProScan search I used to improve the retroviral domain predictions, this unusual prediction is specified by the Phobius (<phobius.sbc.su.se>) protein topology match. 3’-ORF is encoded in the reading frame (+2) and (+1) in IV-8711 in N2(Bristol) and IV-8719 in CB4856(Hawaii) proviruses (Fig.3.F) and (Table.2.) respectively. The 3’-ORF is defined by the F58H7.5 prediction. Predicted 3’-ORF embeds the F58H7.5 starting from position 53 until 324, followed by the in-frame stop codon conserved in both isolates. Provided the F58H7.5 coding sequence is continuous (intronless gene) it is estimated it constitutes approximately 48,5% of the 3’-ORF. Within the F58H7.5 sequence there are two non-synonymous substitutions detected (3’-ORF position 87 H->P and 171 E->K) as divergent between IV-8711 in N2(Bristol) and IV-8719 in CB4856(Hawaii) proviruses (Fig.3.F). 3’-ORF is encoded in positions: (6887-8563) and (6895-8571) in IV-8711 in N2(Bristol) and IV-8719 in CB4856(Hawaii) respectively. Remarkably F58H7.5 is an orphan gene, taxonomically restricted to *C. elegans* lineage. Conducted homology based search analysis indicates F58H7.5 is uniquely present within *C. elegans* species, and therefore absent from other caenorhabdids (including the sister species). This narrow taxonomic distribution of F58H7.5 homology is in the stark contrast with the other proviral frames encoding for the typical retroviral domains described above.

**Table 2.** Open Reading Frames used by predicted protein domains, presented in 5’->3’ order in IV-8711 in N2(Bristol) and IV-8719 in CB4856(Hawaii) proviruses. Discordant frames in the catalytic integrase and 3’-ORF are highlighted by doubly lined frame.* Integrase reading frames were trimmed at N-terminus to limit the mis-alignment due to the frame-shifting mutations (Fig.2.).

| ORF | gag | Pro | Pol | RNaseH | Integrase<br>(162nt) | 3'-ORF | 5'→3' |
| --- | --- | --- | --- | --- | --- | --- | --- |
| Reading<br>frame used | +1 | +1 | +2 | +2 | +3 | +2 | IV-8711<br>(N2) |
|  | +1 | +1 | +2 | +2 | +2 | +1 | IV-8719<br>(Hawaii) |
| Identity(%) | 95/95<br>(100%) | 95/95<br>(100%) | 175/179<br>(98%) | 121/121<br>(100%), | 131/131<br>(100%)* | 557/559<br>(99%) |  |
| similarity(%) | 95/95<br>(100%) | 95/95<br>(100%) | 177/179<br>(99%) | 121/121<br>(100%) | 131/131<br>(100%)* | 558/559<br>(99%) |  |

## Comparison with RETR-1 (III: 8852597 - 8861482)

Provided we demonstrated IV-8711 in N2(Bristol) and IV-8719 in CB4856(Hawaii) proviruses, are ‘identical-by-descent’ and diverged only in non-LTR regions, we compared the isolate specific sequences to RETR-1 retrovirus (Britten 95). RETR-1 is distinctly detected by LTR finder (Table 1), specifically in the N2(Bristol) Chromosome III sequence. This asymmetric display of the element is different from observed for IV-8711 and IV-8719 insertion, apparently present in both isolates. While the description and detailed analysis of the RETR-1 is behind the scope of this work (will be described eswhere), I compared IV-8711 in N2(Bristol) and IV-8719 in CB4856(Hawaii) proviruses to asymmetric insertion of the RETR-1 inactivating the plg-1 gene. RETR-1 (Table. 1) as displayed in N2(Bristol) chromosome III, is 8.886 kb in length (with LTR finder predicted 5’ and 3’ LTR sequences 518 bp and 513 bp respectively). The particular reason of choosing the RETR-1 for the comparative analysis is that in contrast with most other LTR finder predicted retroviral insertions listed with (Table 1.), insertion into chromosome IV shared between N2 and CB4856 aligns significantly with BLAST analysis with relatively well characterized nematode endogenous retrovirus RETR-1. The overall domain architecture of the RETR-1 used for comparison is slightly different to what is described in the previous section concerning the IV-8711 in N2(Bristol) and IV-8719 in CB4856(Hawaii) proviruses. The InterProScan search on the coded, stop masked RETR-1 conceptual translation (described in materials and methods), revealed the all relevant features of the protein domain organization recognized other retroviruses, are grouped into frame (+1). This single (+1) frame of the RETR-1, is fairly long and apparently continuous (uninterrupted by the in-frame stop codons). InterProScan on the RETR-1 single frame (+1) identified regions encoding for conserved protein domains corresponding to reverse-transcriptase polymerase (PS50878/ InterPro:IPR000477), RNaseH (CDD: cd09274), and catalytic Integrase (PS50994/InterPro:IPR001584). In contrast, two additional protein domains typical for the retrovirus genome architecture, identified in the provirus inserted into N2/CB4856 ancestral locus on chromosome IV, was not identified in the RETR-1 using the same search criteria. Notably Pfam protein domain signature matches (PF03732/InterPro:IPR005162) specifying gag gene and (PF13650) specifying retroviral-type Aspartyl Protease gene respectively, are apparently missing from (+1) long reading frame encoded by the RETR-1 (and remaining reading frames). Positively identified protein domain signature matches are compared in the Table.3 (Individual alignments are included in supplementary materials). This analysis demonstrates the 3’-ORF embedding F58H7.5 prediction is a unique and distinct part of the proviral insert in IV-8711 in N2(Bristol) and IV-8719 in CB4856(Hawaii), but consistently absent from the RETR-1 (and any other presently known reverse-transcribing agents (not shown)).

**Table 3.**
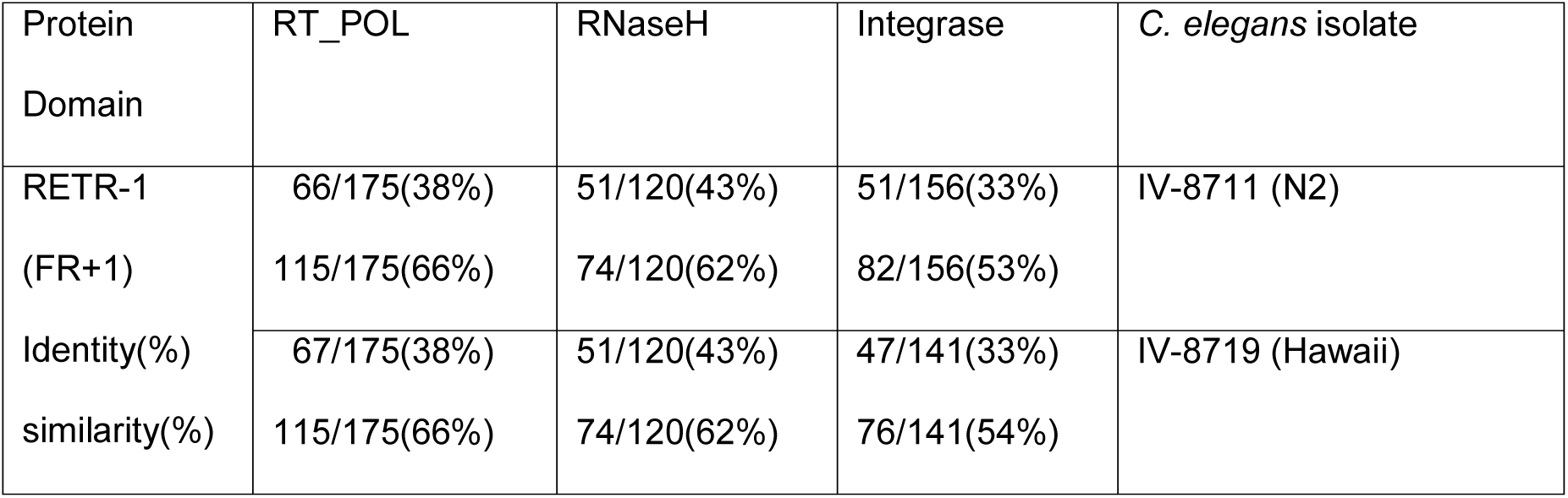
InterProScan identifiable retroviral protein domain signature matches of the retrovirus inserted into chromosome IV-8711 of N2 (reference assembly) and assembled IV-8719 of CB4856 (Thompson et al. 2015) compared to RETR-1 (III).

## Comparison with other reverse-transcribing viruses

Given the original description and sequence analysis described in (Britten 1995) indicated the remarkable homology in the RT polymerase region of RETR-1 to Cauliflower Mosaic Virus (CaMV) RT (Table 1. in (Britten 1995)), we inferred that the insertion detected by LTR finder at the *C. elegans* chromosome IV (IV-8711 in N2 and IV-8719 in CB4856) represent related clan of endogenous retroviruses. The reason for that is the reverse transcriptase polymerase region predicted by the InterProScan in both IV-8711 in N2 and IV-8719 in CB4856 (Fig.3.C), aligns with the CaMV RT and reverse transcriptases of other Caulimoviruses (not shown). Therefore, the reverse transcriptase-polymerase domain phylogenetic tree was build, using the predicted RT protein domains of retroviruses and other reverse-transcribing agents (Fig.4). Both reverse-transcriptase polymerase domain and RNaseH domain fast-tree cladograms of the reverse-transcribing agents, the Caulimoviruses are nested within IV-8711-IV-8719/RETR-1 group. Consistently, with two types of Maximum-Likelihood analysis (where RaxML and PhyML nodes are supported by a bootstrap (Efron et al. 1996)), Caulimoviruses are nested within the IV-8711-IV-8719/RETR-1 group. In particular, the reverse-transcriptase polymerase domain cladograms include the RT-polymerase domain predicted on HBV (Hepatitis B virus), which appear as an outgroup. The reverse-transcriptase polymerase domain cladograms place the retrovirus clade and IV-8711-IV-8719/RETR-1/Caulimovirus branch as sister taxa. The same relation is observed in the cladorograms build based on RNaseH domains, however here the outgroup root (HBV) is removed. Together in all examples provided (catalytic integrase domain tree is not relevant here, as Caulimoviruses in most cases maintain their DNA episomaly (Hull et al. 1987) and thereof do not encode retroviral type integrase) IV-8711-IV-8719/RETR-1 group is basal to Caulimovirus clade and therefore IV-8711-IV-8719/RETR-1 group appears as paraphyletic taxon with respect to pararetroviruses. This result seems suprising as Pararetroviuses existing in plants are currently grouped with Hepadnaviruses (Coffin et al. 1997).

**Figure 4.**
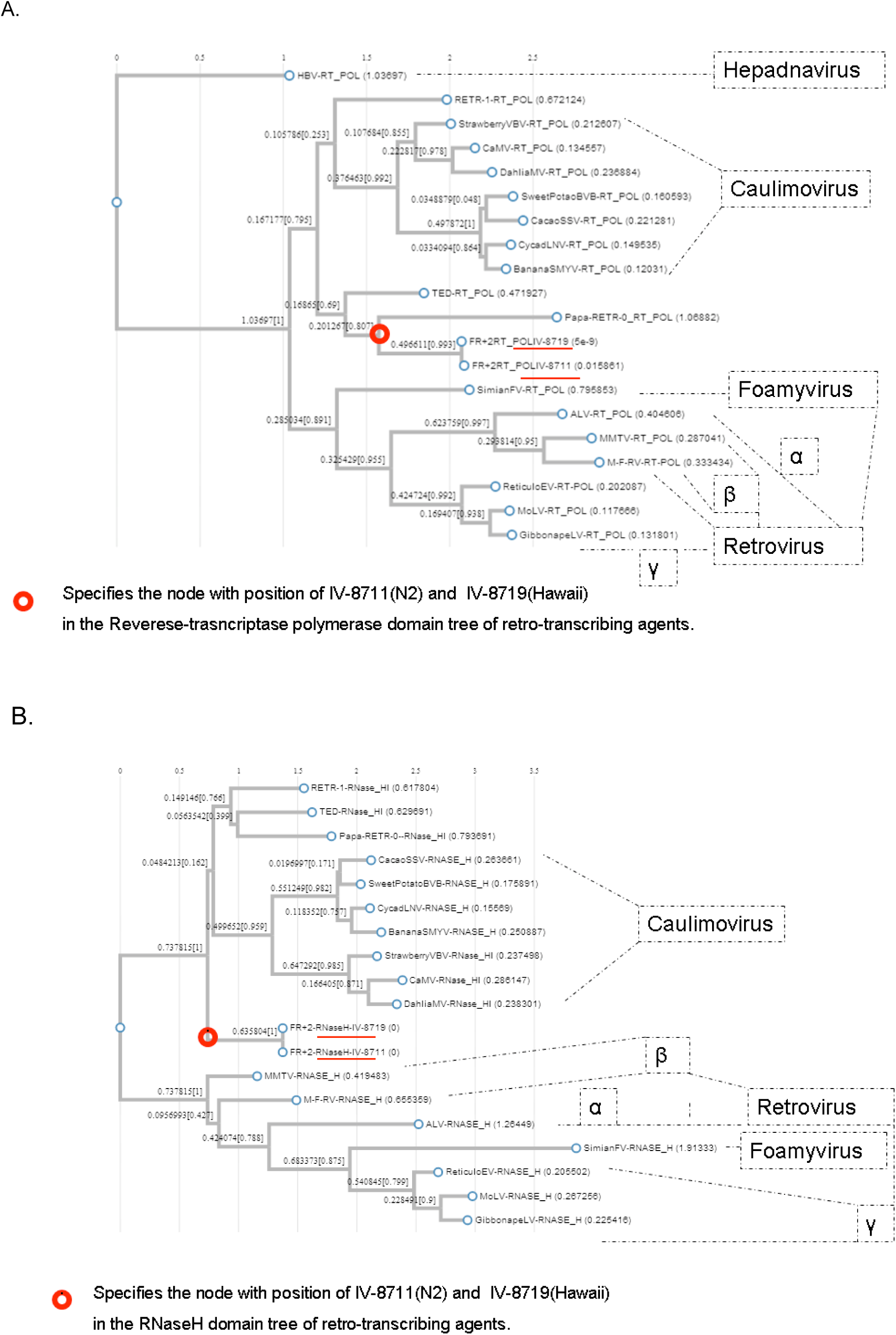
Retroviral protein domain cladograms. Reverse-transcriptase polymerase A. RNaseH B. (description in the text)

## Discussion

Retroviral insertions into animal genomes are established metrics of dating the evolutionary time based on the LTR divergence. According to the retrovirus replication model (Coffin et al. 1997) accepted for the endogenous retroviruses and the LTR-retrotransposons, the LTRs are identical at the time of an insertion, than subsequently diverge, acquiring independent mutations at neutral rate. Examples of dating of the retroviral insertions and estimates of the evolutionary splits are available in the literature i.e. (Johnson and Coffin 1999) (Yohn et al. 2005) (Arnaud et al. 2007) (Jo et al. 2012) and are based on the calculations which in principle could be used and applied towards dating of retroviral insertions in *C. elegans* isolates. Those estimates could be useful in dating of the evolutionary split between the reference N2(Bristol) and Hawaiian isolates. In a given example, the conserved insertion pair of the IV-8711 in N2 and IV-8719 in CB4856, diverged only minimally. Presented evidence concerning the divergence in the LTR region indicates the long terminal repeats in both isolates did not acquired any new mutations since a split from the prototype ancestral *C. elegans*. Therefore, I regard IV-8711 insertion represents the evolutionary recent event. According to the accepted model this suggests the split between N2(Bristol) and Hawaiian isolates could be estimated as remarkably recent, essentially behind the resolution power of the dating. On the other hand however, within the presented set of the retroviral insertions conserved between N2(Bristol) and Hawaiian isolates, there are examples which contradict the above evolutionary scaling (data not shown). In few examples there is an evidence for LTR acquired mutations prior to split of N2(Bristol) and Hawaiian isolates. In those examples, the 5’- and 3’-LTRs diverged but the particular substitution is same in both isolates. The presence of this class of substitutions conserved in both isolates, suggest some of the insertions into prototype *C. elegans* genome have persisted for sufficient time in the ancestral *C. elegans* line to permit for the LTR divergence prior to split of an N2 and CB4856. Provided the alternative estimates for the split of N2 and Hawaii have been proposed (Thomas et al. 2015), I suggest those might need to be revised or rescaled using the metrics based on the LTR divergence. While detailed follow up analysis on those particular examples is emerging, it however remains behind the scope of this description.

The architecture of the unusual 3’-ORF embedding the F58H7.5 is described in the results section. F58H7.5 is an uncharacterized *C. elegans* gene of presently unknown function encoding for an novel protein. Based on the homology searches it seems the F58H7.5 is a distinct and unique, taxon restricted, species specific, orphan gene. The above orphan gene assignment is for the following rationale: F58H7.5 predicted protein is not associated with any of the known protein families represented in *C. elegans* neither elsewhere in the taxonomy tree. Homology based search on predicted protein F58H7.5 indicates no informative homologies in other caenorhabdids neither the sister species *C.briggsae*^1^. Moreover, no additional homology is found in searches excluding the *Caenorhabditis* genus. The F58H7.5 gene is clearly embedded into provirus prediction on chromosome IV-8711 in N2(Bristol) and IV-8719 in CB4856(Hawaii) and is predicted to lay in the region between the catalytic integrase and 3’ LTR (result section and Fig.1.C). The F58H7.5 gene is encoded in single frame in continuous, uninterrupted manner and is apparently expressed (by virtue of the annotated mRNA (reference sequence # NM_067613.1 ║ GI:17540863)). Lack of introns spanning the coding sequence appears markedly relevant, as is consistent with the retroviral origins of F58H7.5. Retroviruses and other reverse-transcribing agents, do replicate via RNA intermediate (Baltimore 1970) and (Temin and Mizutani 1970) and it is widely accepted that introns inserted in the sense (+) orientation will be removed from genomic transcripts (Boeke et al. 1985) prior to insertion into chromosomal loci. Provided the above rationale, as main conclusion of the described work, I suggest the annotated gene F58H7.5 in the reference N2 assembly, is a retroviral-borne gene acquired via RNA intermediate, specifically into *C. elegans* linage prior to the split of the N2(Bristol) and CB4856(Hawaii) isolates. I note further, that while it has long been established the retroviruses could embed the non-viral genes (or its portions) and insert those into chromosomal loci (Spector et al. 1978), the acquisition of the taxon restricted, species specific gene however has not been previously postulated.

The insertion of the retrovirus presently located on chromosome IV-8711 in N2 and IV-8719 in CB4856, represent the example of the ancestral retroviral infection contrasting the RETR-1 insertion on chromosome III. While both IV-8711/IV-8719 and RETR-1 share the protein domains typical for the organization of endogenous retrovirus genome, the RETR-1 display in the LTR finder analysis is polar i.e. RETR-1 is detected explicitly in N2(Bristol) but missing from CB4856(Hawaii) (Table.1. and author’s analysis. Not shown.). RETR-1 was identified by *C. elegans* genome sequence analysis (Sulson and Waterston 1998) and is inserted into chromosome III of the reference N2 (spanning the clone border F44E2/PAR3 Alan Coulson pers communication 1997). In the N2 reference sequence the RETR-1 is regarded transcriptionaly active in germline and embryonic tissues, as judged by multiple ESTs sequences and *in situ* hybridization (Yuji Kohara, personal communication 1997, and NEXTdb pattern <nematode.lab.nig.ac.jp/db2/ShowCloneInfo.php?clone=88f8>). Others (Maydan et al. 2007 and Maydan et al. 2010) have found by comparative genomic hybridization that N2 RETR-1 locus is polymorphic (with large deletion spanning the RETR-1 insertion site in N2) when compared to CB4856. Consistently, (Papoli et al. 2008) demonstrated that RETR-1 is inserted into F44E2.11 in N2 and concluded is absent from plg-1(+) CB4856. Those findings are in the substantial agreement with the results presented here (Table 1.). First RETR-1 is accurately predicted by the LTR finder and served as positive control in the genome-scale search. Second major retroviral protein domains predicted in the stop-masked frames of the ancestral retroviral element inserted on chromosome IV, appear conserved with domains predicted on the RETR-1, including reverse-transcriptase, RNaseH and catalytic integrase. The presented results however demonstrate, that comparing the gag gene region predicted consistently on the stop-masked frame (+1) the IV-8711 in N2 and IV-8719 in CB4856 is absent from RETR-1 (by direct comparison and by *de novo* InterProScan analysis). This rise the possibility that gag gene in the RETR-1 (III) element is either rearranged (deleted) or diverged significantly from other members of the clade, to sufficient degree to prevent detection. Indeed the rearrangements appear common among endogenous retroviral insertions (authors unpublished observation). In this perspective the RETR-1 often regarded as ‘active’ (the original communication of (Britten 1995) but also by (Dennis et al. 2012) who tackled the RETR-1 experimentally), might in the fact represent immobile (albeit transcriptionaly active) insertion into plg-1 gene. Third, considering the evolutionary scale since the insertion of into the locus on chromosome IV present in both N2 and Hawaii isolates and RETR-1, it appears the later element was inserted into the N2 genome after the split of the Hawaii from the ancestral *C. elegans* line. While this model would possibly explain the polar distribution of the RETR-1 (and possibly some other elements detected discretely in the N2 (Table.1.) but not in CB4856), the results of the LTR alignments appear to contradict this perspective. As mentioned above the 5’ and 3’ LTRs in the elements present in the IV-8711 in N2 and IV-8719 in CB4856, are identical. In contrast the 5’ and 3’ LTRs predicted by the LTR finder on the RETR-1 are different in length (518 and 513 bp respectively (Table 1.), suggesting the insertion or deletion must have occurred) and clearly acquired aberrant bases (>10 aberrant bases between 5’ and 3’ LTR) at the 3’ end, since the time insertion. Importantly, the LTR finder predicts so called TSR (Target Site Repeats, short duplicated sequences of the host at the integration site), implicated to occur by the accepted retroviral integration model (Brown 1997) adjacent to RETR-1 insertion site [TSR: CGACTA]. The presence of the TSRs flanking the RETR-1 insertion site might rule-out the inaccurate prediction of LTRs. Considering, the observed divergence rate of the LTRs in the RETR-1 and the divergence rate of the LTRs present in IV-8711 in N2 and in IV-8719 in CB4856, those findings appear contradictory, if the neutral substitution model of the LTRs divergence is assumed. This inconsistency might be due to selective pressure on some endogenous retroviral insertions loci (i.e. serving as the host restriction factors) or the effects of the negative selection (i.e. if the insertion results in the trait affecting the overall fitness of the carrier) or perhaps for some other reasons. This note however should serve as the word of caution, considering the long distance evolutionary extrapolations based on the single isolated example (given the examples supporting other evolutionary scenarios could be drawn out of the examples given in (Table.1.). Those examples will be discussed elsewhere).

Phylogenetic reconstructions of the predicted retroviral protein domains. The intimate kinship established for the ancestral *C. elegans* provirus IV-8711 in N2 and in IV-8719 in CB4856 and RETR-1(III) is further supported by the phylogenetic reconstructions. Based on the reverse-trascriptase polymerase domain cladograms of the retro-transcribing viral agents, IV-8711-IV-8719/RETR-1 like elements appear as paraphyletic group with Caulimovirus clade. Cladograms establish the sister relationship between retrovirus clade and caulimovirus/IV-8711-IV-8719/RETR-1. The above grouping is supported by placement of the RT-polymerase domain predicted independently in genomes of seven different Caulimoviruses and seven different retroviruses (representing major branches in animal retrovirus taxonomy) respectively, in addition to five members of the IV-8711-IV-8719/RETR-1 group. It appears even more striking as the same phylogenetic kinship is supported by the RNaseH domain cladograms. [In contrast to cladograms build based on RT-polymerase domain (when single defined domain PS50878/ InterPro:IPR000477 is predicted on all genomes included into tree) RNaseH domain is predicted either as RNaseH (PS50879 RNASE_H domain, found typically in the vertebrate retroviruses) or RNaseHI (cd09274 RNase_HI_RT_Ty3 found in the invertebrate retroviruses including the group of IV-8711-IV-8719/RETR-1), while both types of the RNaseH domain are identified in Caulimoviruses. Provided the IV-8711-IV-8719/RETR-1 group contains elements infecting nematodes and insects (and other retroviruses that are associated with invertebrates other than insects and nematodes, unpublished observation) the grouping with Caulimoviruses is surprising given the later are found exclusively infecting plants. Curiously, the association between described provirus inserted into chromosome IV and Caulimoviruses, extends beyond the homology found in the polymerase domain. Insertion into chromosome IV-8711 in N2 and IV-8719 is predicted by the LTR finder to utilize the methionyl-tRNA as a primer for reverse transcription reaction. This coincides with the tRNA preference described for Caulimoviruses (Hull et al. 1987), despite the overall differences in the genome replication via reverse transcription (however this Methionyl-tRNA bias remains speculative, as is based solely on the computational prediction and thereof requires the experimental verification). Regarding the above findings in the perspective concerning the scope of this description, it needs to be emphasized, that while the above phylogenetic reconstructions robustly confirm the identified region of the *C.elegans* genome represents the inserted endogenous retrovirus, the phylogenetic tools can not be applied efficiently towards the identified 3’-ORF embedding the F58H7.5 gene. This is because (as noted above) the F58H7.5 is an orphan gene, taxonomically restricted to *C. elegans* lineage, precluding the cross-taxa comparisons. The variant of the F58H7.5 identified in CB4856 (result section Fig. 3.F) is than the first known and constitutes the only prediction available for the phylogenetic comparisons.

Considering the presented results in the genome-scale perspective, the work focuses on the single example of the past infection of the *C. elegans* by the retrovirus, represented as an insertion on the proximal arm of the chromosome IV present in two modern geographical isolates. As mentioned in the initial sentence of the results section, the detailed results of the genome scale analysis are out of the scope of this description and will be described elsewhere. *C. elegans* genome was analyzed with the LTR finder program and the combined results are outlined in (Fig.5) and summarized in (Table 4.). In total LTR finder predicted 567 and 588 candidate loci in the N2(Bristol) and CB4856(Hawaii) respectively. Of above only fourteen predictions are listed in (Table 1.) where thirteen was verified as retroviral insertions. Would there be many more endogenous retrovirus insertions demanding the description? While genome-scale searches in *C. elegans* for the endogenous retroviruses, where attempted by others (reviewed in (Bessereau 2006)) for various reasons, the entire picture remains blurred. There is an instructive example included into (Table 1.) of another provirus inherited from the ancestral *C. elegans*: the X inserted element X-12435. This particular element is predicted on both N2 and CB4856 genomes by the LTR finder by the reciprocal LTR match. However, in given case LTR finder internally enabled ScanProsite detects the retroviral domain only on the proviral locus in N2(Bristol) chromosome X but ignores the domain encoded by the orthologous locus on CB4856(Hawaii) X. While the reasons for this behavior remains largely enigmatic (i.e. it is formally possible that the domain encoding sequences deteriorated enough to prevent the detection by the internally enabled ScanProsite) this example suggests, there might be other proviral insertions where support by the internally enabled ScanProsite is insufficient to detect the protein domains supporting the search. To compensate for that insufficiency, I propose, the frame-wise protein domain search would be enabled externally on the conceptually translated, coded and stop-masked amino-acid strings. The presented evidence implies, the implemented stop-masking offers the improvement detecting the endogenous retroviruses insertions with InterProScan search coupled with the LTR finder output. I further suggest, the stop-masking approach implemented here, could be easily generalized into genome scale analysis, with the following assumption: i. sequence of the entire chromosomes is frame-wise translated into amino-acid coded strings, ii. coded amino-acid strings are than stop-masked and searched for the co-occurring adjacent retroviral domains indicative for the insertion of the provirus iii. the possible LTR pairs are verified by the LTR finder.

## Footnotes

1 The WormBase homology information specifies the weak (BLAST e-value 5.8e-06 over the 31.2% of the protein length) match with CBG12079 gene. CBG12079 (119 amino-acid residues, were 69 residues is a repeat of continued stretch of homo-polymeric charged glutamates, below) is considerably smaller gene, compared to F58H7.5 (272 amino-acid residues), and is interrupted by single intron. CBG12079 gene itself is found in ~3.5kb intron of Cbr-unc-73 on Cb-Chr-I, and therefore collectively unlikely orthologous with F58H7.5. [available via http://www.wormbase.org/species/c_briggsae/protein/BP:CBP22639#06--10> >CBG12079 MISHRGIVKFDTINSQFLWLSSTSKRACQSISSTDEKEEEEEEEEEEEEEEEEEEEEEEEEEEEEEEEEEEE EEEEEEEEEEEEEEEEEEEEEEEEEEEEEEEEEEEQKMRIRWMWITK)]

